# Genetic Diversity and Structural Complexity of the Killer-Cell Immunoglobulin-Like Receptor Gene Complex: A Comprehensive Analysis Using Human Pangenome Assemblies

**DOI:** 10.1101/2023.11.12.566753

**Authors:** Tsung-Kai Hung, Wan-Chi Liu, Sheng-Kai Lai, Hui-Wen Chuang, Yi-Che Lee, Hong-Ye Lin, Chia-Lang Hsu, Chien-Yu Chen, Ya-Chien Yang, Jacob Shujui Hsu, Pei-Lung Chen

## Abstract

The killer-cell immunoglobulin-like receptor (KIR) gene complex, a highly polymorphic region of the human genome that encodes proteins involved in immune responses, poses strong challenges in genotyping due to its remarkable genetic diversity and structural intricacy. Accurate analysis of KIR alleles, including their structural variations, is crucial for understanding their roles in various immune responses. Leveraging the high-quality genome assemblies from the Human Pangenome Reference Consortium (HPRC), we present a novel bioinformatic tool, the Structural KIR annoTator (SKIRT), to investigate gene diversity and facilitate precise KIR allele analysis. We applied SKIRT on 47 HPRC-phased assemblies and identified a recurrent novel *KIR2DS4/3DL1* fusion gene in the paternal haplotype of HG02630 and maternal haplotype of NA19240. Additionally, SKIRT accurately identifies eight structural variants and 17 novel nonsynonymous alleles, all of which were independently validated using short-read data or quantitative polymerase chain reaction. Our study has discovered a total of 570 novel alleles, among which eight haplotypes harbor at least one KIR gene duplication, six haplotypes have lost at least one framework gene, and 75 out of 94 haplotypes (79.8%) carry at least five novel alleles, thus confirming KIR genetic diversity. These findings are pivotal in providing insights into KIR gene diversity and serve as a solid foundation for understanding the functional consequences of KIR structural variations. High-resolution genome assemblies offer unprecedented opportunities to explore polymorphic regions that are challenging to investigate using short-read sequencing methods. The SKIRT pipeline emerges as a highly efficient tool, enabling the comprehensive detection of the complete spectrum of KIR alleles within human genome assemblies.

## Introduction

Killer-cell immunoglobulin-like receptors (KIR), expressed primarily on the surface of natural killer (NK) cells and specific T lymphocytes, are essential and epistatic for interacting with human leukocyte antigens (HLA) on target cells to regulate host innate and adaptive immunity (Martin et al. 2002; Vilches and Parham 2002; Pollock et al. 2022), modulating the activation and inhibition of NK cell cytotoxicity and cytokine production. The KIR complex comprises 17 KIR genes with diverse inhibitory and activating domains, although not all are essential for healthy individuals. KIR haplotypes exhibit extensive diversity through expansion and recombination (Shilling et al. 1998), with gene contents ranging from eight to 14 KIR genes (Hsu et al. 2002). The Genome in a Bottle (GIAB) consortium excluded KIR regions from the benchmarking regions (v4.2.1) because of their copy number variability (Wagner et al. 2022), highlighting the complexity of KIR genotyping. The accurate determination of KIR gene copy numbers and high-resolution allelic genotypes can substantially facilitate research on therapeutic pathways for immune-related disorders (Béziat et al. 2013).

The biological and clinical significance of KIR genes is rooted in their involvement in various immune-mediated diseases and their impact on hematopoietic stem cell and organ transplantation (Boudreau et al. 2017; Zamir et al. 2022; Dizaji et al. 2021). Associations have been reported between specific KIRs or KIR-HLA combinations and autoimmune diseases such as rheumatoid arthritis, psoriasis, type 1 diabetes, and reproductive health issues (Alexandrova et al. 2022; Feng et al. 2022). KIR genes have also been implicated in infectious diseases, including human immunodeficiency virus (HIV)and hepatitis C virus (HCV) infections, where specific KIR and HLA combinations have been shown to influence disease progression and the response to antiviral therapy (Khakoo and Carrington, 2006; Boudreau et al. 2016; Boelen et al. 2018; Li et al. 2022). Moreover, KIR gene diversity has been linked to susceptibility and clinical course of human cancers, including leukemia, lymphoma, and melanoma (Pende et al. 2019). Clinical trials, *in vitro* and *in vivo* studies have shown promising results using a combination of lirilumab, an anti-KIR monoclonal antibody (mAb) that targets the interactions between *HLA-C* and *KIR2DL1/L2/L3*, and anti-PD-1/PD-L1 immunotherapies for the treatment of squamous cell carcinoma of the head and neck (SCCHN) and human papillomavirus (HPV)-positive cervical cancer (Hanna et al. 2022; Liu et al. 2022). The results of another phase I/II clinical trial of lirilumab combined with an anti-PD-1 mAb in patients with multiple myeloma are also awaited (Clara and Childs 2022). The multifaceted functions of KIRs and their combination with HLA stem from their expression during the development of NK and T cells (Bashirova et al. 2006; Fauriat et al. 2010). Its expression is influenced by both copy number variations (CNVs) and allelic polymorphisms at KIR gene loci, which are shaped by sequence variations in the coding, promoter, and other non-coding regions (Yawata et al. 2006). These variations can modulate factors, such as binding affinity, specificity, ligation angle and orientation, signal transduction capability, and intracellular trafficking between receptors and their cognate ligands, contributing significantly to the diversity of KIR-HLA interactions (Gardiner et al. 2001; Saunders et al 2016; Wu et al. 2021).

KIR regions are usually characterized as A and B haplotypes, both of which contain four framework genes: *KIR3DL3*, *KIR3DP1*, *KIR2DL4*, and *KIR3DL2*. Haplotype A is the majority worldwide, with homozygous AA predominant (de Brito Vargas et al. 2021). However, many unknown haplotypic structures, gene contents, or receptors with novel functions may exist, but remain challenging to identify using current methods. Previous studies have revealed CNV haplotypes in multiple genes such as *KIR3DP1*, *KIR2DL4*, and *KIR3DL1/S1* (Williams et al. 2003; Gómez-Lozano et al. 2005). Duplications and deletions of *KIR3DP1*, *KIR2DL4*, and *KIR3DL1/S1* appear to form a highly conserved triad with high frequency in the same type of unequal crossover events throughout human evolution (Martin et al. 2003; Williams et al. 2003; Amorim et al. 2021). However, many studies have overlooked the CNVs of framework genes, and most KIR genotyping algorithms presume a diploid genome with two copies of each framework gene. Based on this assumption, they calculated the copy numbers of all other KIR genes, potentially leading to limited or incorrect results (Norman et al. 2016; Marin et al. 2021; Sakaue et al. 2022; Song et al. 2023). Traditional methods require manual crafting of haplotypes by incorporating gene content and copy number information, followed by estimation based on the linkage disequilibrium (LD) of KIR genes and their alleles derived from known KIR haplotypes (Amorim et al. 2021). The absence of correct phasing information complicates the investigation of structural variations among KIR genes, including crossover events, gene deletions, and gene duplications. These challenges are compounded by the ongoing expansion and evolution of KIR genes in all populations (Alicata et al. 2020; NurWaliyuddin et al. 2022; Kevin-Tey et al. 2023).

Kass assembly method was introduced as an efficient approach for interpreting KIR diploid haplotypes by assembling contigs from probe-captured PacBio CCS long-read sequencing data (Roe et al. 2020). Assembly quality, including completeness and contiguity, largely depends on the sensitivity and specificity of the in vitro and in silico probes targeting the highly polymorphic KIR region, and the assembly algorithm employed. Therein lies the potential scope for refinement to further enhance output quality. The Telomere-to-Telomere (T2T) consortium achieved a significant scientific milestone by successfully sequencing the complete haploid human genome, T2T-CHM13, which encompasses contiguous sequences for each autosome and X chromosome (Nurk et al. 2022). Later, the Human Pangenome Reference Consortium (HPRC) released forty-seven high-quality, haplotype-resolved genome assemblies (HPRC-release) (Wang et al. 2022; Liao et al. 2023). These were created using the graph trio binning algorithm, Hifiasm (Cheng et al. 2021), and family trio samples from the 1000 Genomes Project (1KGP) (The 1000 Genomes Project Consortium 2015). The assembly of HG002, one of the 47 genomes, was characterized by more accurate variant calls across single nucleotide variants (SNVs), small insertions/deletions (indels), and structural variants than the HG002 assemblies created using other algorithms (Jarvis et al. 2022). Compared to T2T-CHM13, the paternal and maternal assemblies of HPRC autosomes cover approximately 92.8% and 94.1% of T2T-CHM13, respectively (Liao et al. 2023). The reliability of these 47 HPRC assemblies was estimated to be 99.12%, whereas T2T-CHM13 was designated with a 99.91% reliability score (Liao et al. 2023). This level of accuracy, near-complete coverage, and almost telomere-to-telomere continuity underscores the value of the 47 HPRC diverse diploid genome assemblies. Furthermore, they facilitate the study of long, complex, and repeat-rich regions of the genome, such as the major histocompatibility complex (MHC) and KIR.

The Kass annotation workflow effectively annotates the human genome assembly generated by its assembly workflow with high-resolution allelic genotyping information for KIR genes (Roe et al. 2020). However, some erroneous annotations were generated when tested with the same HPRC-released HG002 data (Supplemental Table 1), suggesting a need for improved methods for the high-resolution annotation of haplotype-resolved assemblies, particularly in the context of CNVs in framework genes. On the other hand, T1K (Song et al. 2023) also utilized 26 of the 47 HPRC genome assemblies to benchmark its KIR genotyping algorithm. Nevertheless, the results did not consider the probability of having novel structural variations in the 52 haplotypes analyzed. Therefore, achieving complete high-resolution allelic KIR genotyping of the diploid human KIR region requires a refined method for annotating the human genome assembly with KIR alleles.

In the present study, we developed an improved <u>S</u>tructural <u>KIR</u> anno<u>T</u>ator (SKIRT) method for the high-resolution allelic and CNVs identification of KIR genes. This study aimed to identify and annotate the complete haplotypes of the KIR complex with high-resolution allelic genotypes of all existing KIR alleles, describing the gene copy numbers and structures of all haplotype-resolved personal genome assemblies. Accurate allelic genotyping annotation of the KIR complex enables researchers to precisely examine the haplotype structures, copy numbers, and various structural variations in each phased haploid genome. Our study on long-read assembled genome data confirmed the exact flanking positions, providing accurate alleles of up to seven digits and an accurate copy number of each gene in the KIR region. Complete haplotypes of the KIR complex are crucial for advancing our understanding of the roles of KIR genes in various immune-mediated diseases and their therapeutic implications.

## Results

### Method Overview

The 47 HPRC diploid genome assemblies, produced using Hifiasm with PacBio HiFi reads were processed using our proprietary SKIRT pipeline. This pipeline was engineered to yield the highest possible resolution of KIR alleles (Fig. 1a). Initially, it identified the coding regions of all KIR genes in the assembly contigs and distinguished five-digit alleles by mapping them with IPD-KIR CDS-only sequences. Upon perfect alignment with five-digit alleles, seven-digit alleles were further identified using IPD-KIR genomic sequences. However, when five-digit alleles aligned with variations, a synonymous test was conducted to determine whether these variations led to amino acid alterations, subsequently resulting in either synonymous (three-digit resolution) or entirely novel transcripts (Fig. 1b).

**Figure 1.**
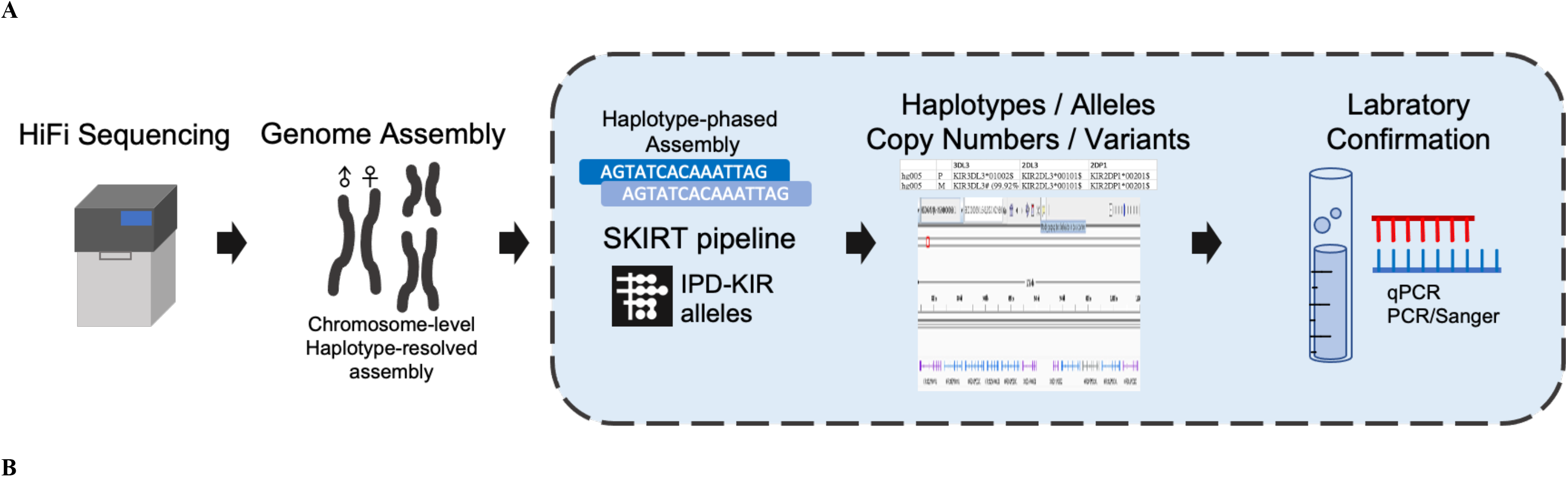
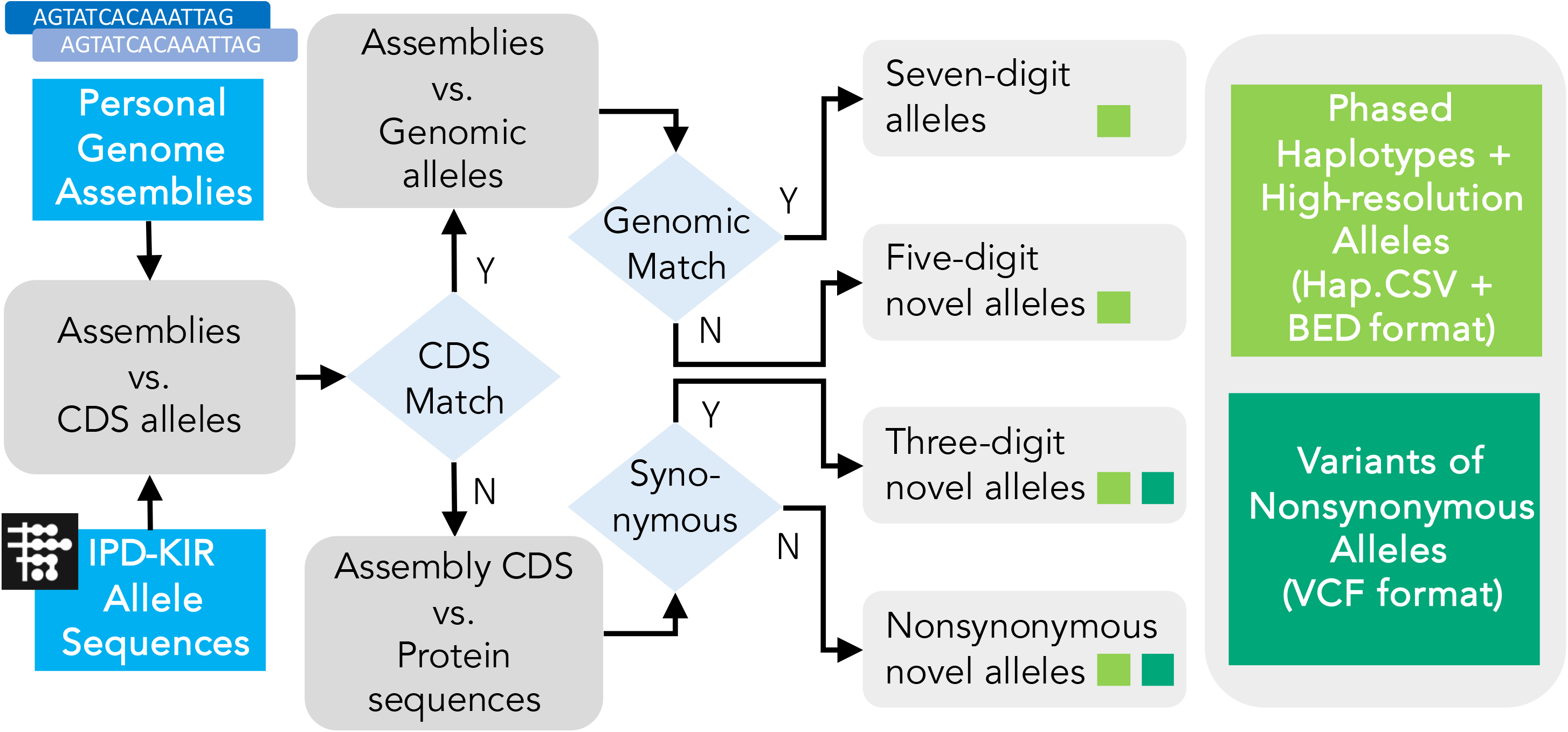
KIR allele annotation of the diploid genome assemblies. **(a) Overview of the high-resolution allelic KIR identification workflow.** The segment shaded blue denotes the primary focus of this study. We developed a specialized pipeline, the <u>S</u>tructural <u>KIR</u> anno<u>T</u>ator (SKIRT), to facilitate annotation of high-resolution KIR alleles and structural variations in human genome assemblies. A series of laboratory experiments were conducted to validate the computational findings and ensure their accuracy. **(b) The operational flow of the SKIRT pipeline.** This pipeline merges a long sequence alignment tool, minimap2 (Li 2018), with in-house Python and Shell scripts. These scripts map the KIR Repository of Immuno Polymorphism Database (IPD-KIR) allele sequences against genome assemblies, such as the HPRC Year-1 released data, to identify existing KIR genes on each contig with an accuracy of up to seven-digit high-resolution alleles.

For nonsynonymous alleles, we conducted further verification using Illumina whole-genome short-read sequences (WGS) and, when necessary, PacBio HiFi reads to authenticate the presence of gene sequences. When the variant percentage surpassed 1%, we further examined the likelihood of novel structural variations such as gene fusion and segmental deletion. Any structural variations identified in our study, including CNVs, deletions, and gene fusion, were further confirmed through a series of laboratory experiments.

### Overview of the 94 HPRC KIR haplotypes

We fully characterized the high-resolution alleles of KIR genes for all known KIR alleles in 47 HPRC human genome assemblies (Figs. 2a-d and Supplemental Table 2). Sixty-three haplotypes (67.02%) were centromeric A and telomeric A (Fig. 2a), seven (7.45%) were centromeric A and telomeric B (Fig. 2b), 14 (14.89%) were centromeric B and telomeric A (Fig. 2c), and 10 (10.64%) were centromeric B and telomeric B (Fig. 2d). Among the 94 haplotypes of the 47 genome assemblies, eight (8.51%) displayed at least one KIR gene duplication, six (6.38%) exhibited deletions of both *KIR3DP1*/*KIR2DL4* framework genes (Figs. 2b,d), and one haplotype showed *KIR3DL2* framework gene deletion (Fig. 2a). In total, 15 haplotypes (15.96%) showed structural variations in KIR genes.

**Figure 2.**
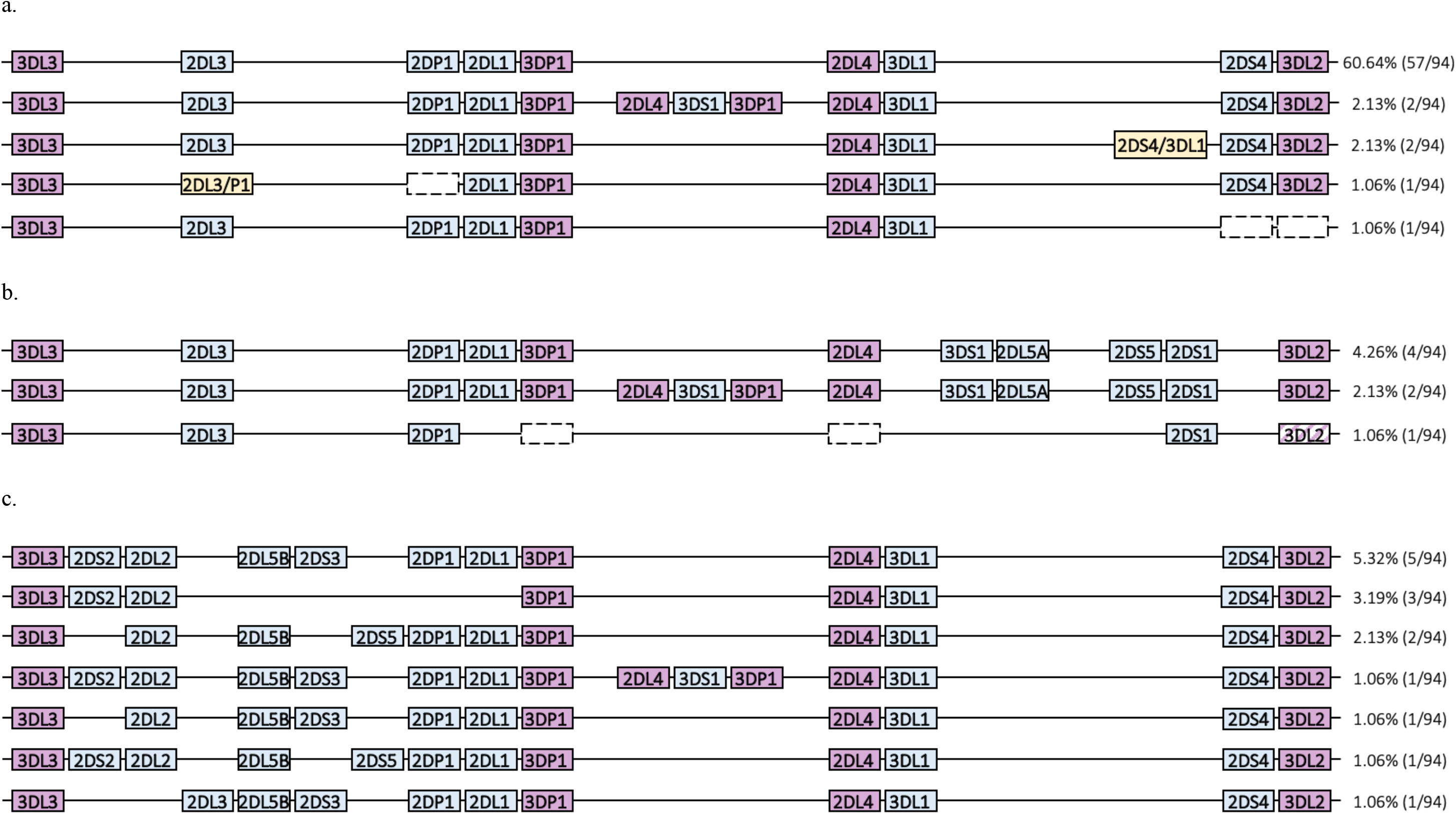
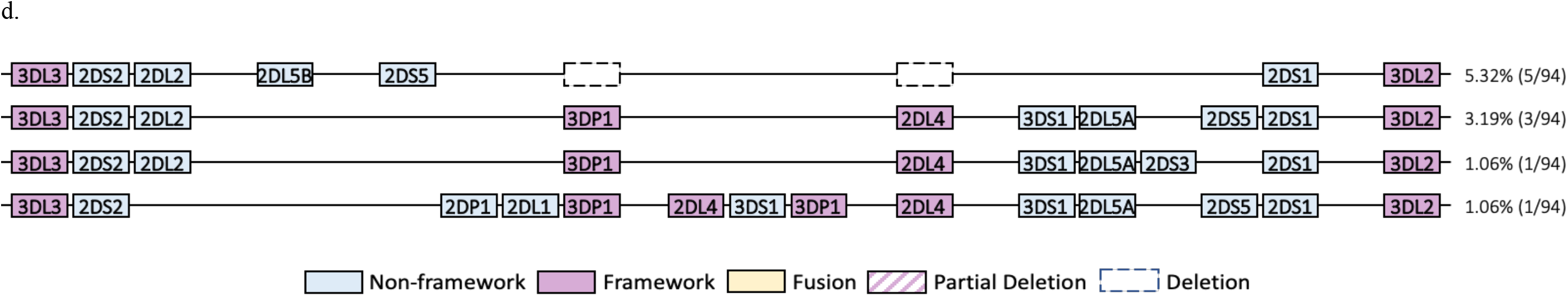
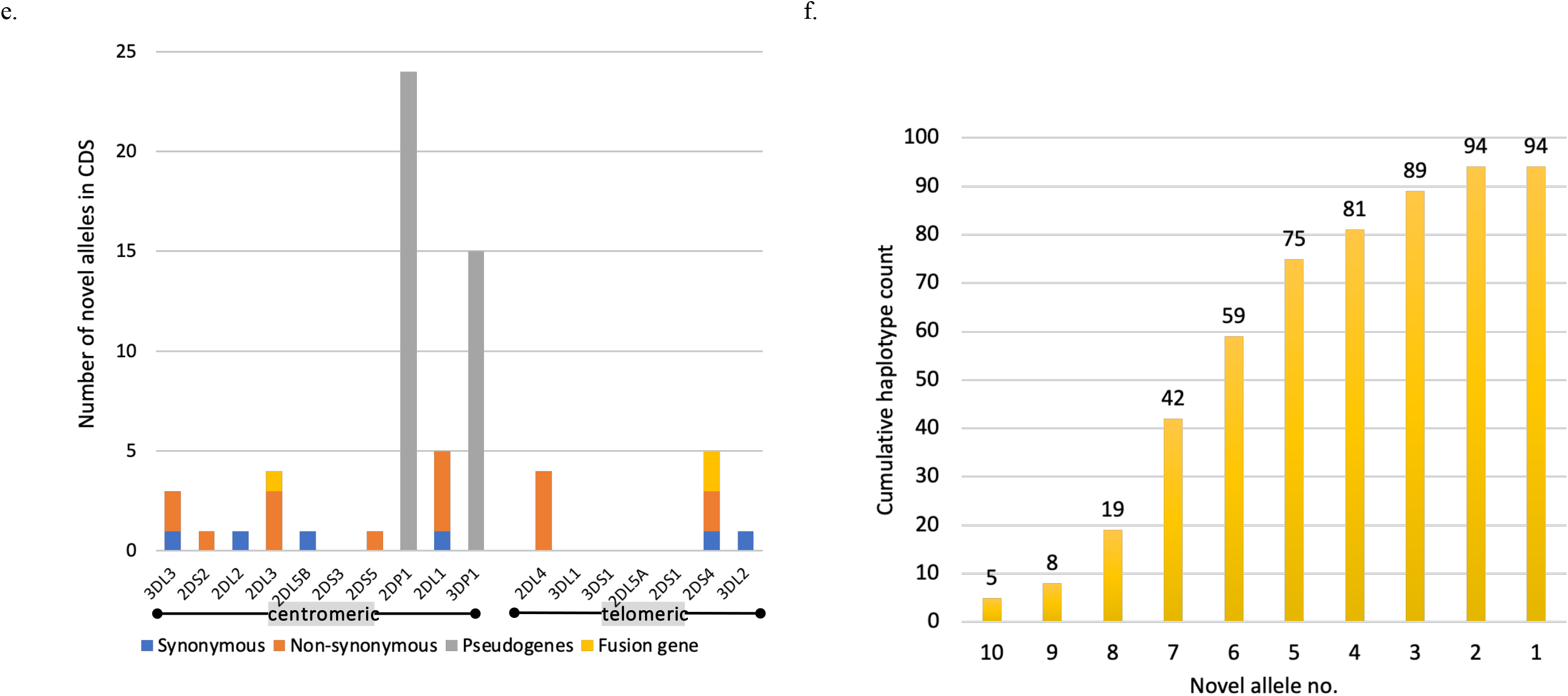
KIR haplotypes and novel alleles in the 47 HPRC phased assemblies characterized by SKIRT. **(a)-(d) Diverse KIR haplotypes and structural variations in the HPRC cohort.** The KIR haplotypes were categorized as **(a)** cA-tA haplotypes (67.02%, 63/94), **(b)** cA-tB haplotypes (7.45%, 7/94), **(c)** cB-tA haplotypes (14.89%, 14/94), and **(d)** cB-tB haplotypes (10.64%, 10/94). **(e) The identified novel alleles with variances in the CDS of KIR genes.** Blue indicates alleles with synonymous variants, whereas orange indicates alleles with nonsynonymous variants. Counts of 2DS3 and 2DS5 were observed in the centromeric region. The pie chart only counts novel alleles of functional KIR genes, excluding fusion genes. **(f) Haplotype counts with novel alleles identified.** The number in each bar indicates the cumulative number of haplotype counts containing at least the number of novel alleles specified on the X-axis. There were five haplotypes with ten novel alleles and all 94 haplotypes contained at least two novel alleles.

In this study, we used the notation SampleID-P to denote the paternal haplotype of the sample and SampleID-M to denote the maternal haplotype, with SampleID representing the sample identity. For example, HG002-P and HG002-M (Fig. 3a) correspond to the paternal and maternal haplotypes of HG002.

**Figure 3.**
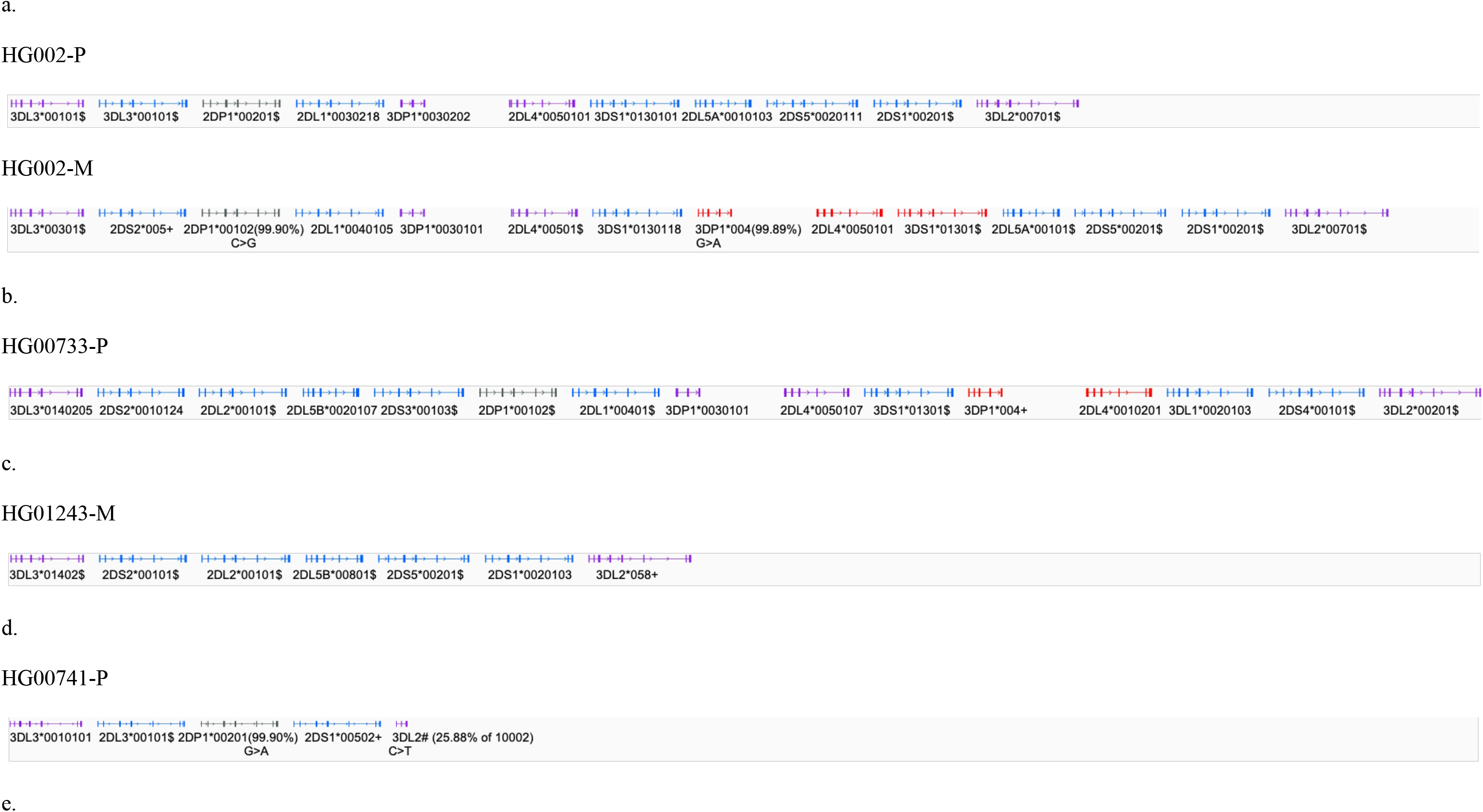
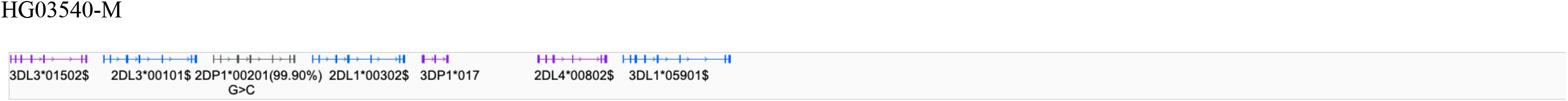
Representation of Typical and Atypical Haplotype Compositions within the HPRC Cohort. (a) HG002 paternal (HG002-P) and maternal (HG002-M) haplotypes. The paternal haplotype of HG002, a typical cA01-tB01 haplotype, is presented from the centromeric to the telomeric motifs without deletions, duplications, ambiguities, and other structural variations. (b) The paternal haplotype of HG00733 features a duplication of *KIR3DP1-KIR2DL4-KIR3DL1/S1*, integrated into a cB01-tA01 haplotype. (c) The maternal haplotype of HG01243 is distinguished by the absence of both the *KIR3DP1* and *KIR2DL4* framework genes. (d) The paternal haplotype of HG00741 is characterized by the deletion of both the *KIR3DP1* and *KIR2DL4* framework genes as well as a truncated *KIR3DL2* gene containing only exons 1–3. (e) The maternal haplotype of HG03540 is marked by a deletion of *KIR3DL2* framework gene with an African high-frequency *KIR3DL1* allele resulting from a deletion between *KIR3DL1* and *KIR3DL2* (Color codes: Purple represents framework genes, red represents duplicated genes, blue represents non-framework genes, and gray represents pseudogenes. Allele suffix marks: #, an allele having nonsynonymous variant(s) in CDS, segmental deletion or fusion with another gene; =, an allele having synonymous variant(s) in CDS; $, an allele having variant(s) in non-CDS; +, a genomic allele matching an IPD-KIR CDS-only allele).

Within the HPRC assemblies, we identified a recurrent duplication event in the *KIR3DP1*-*KIR2DL4*-*KIR3DL1*/*S1* triad in six samples. We also detected a known fusion gene, *KIR2DL3/2DP1*, in HG02723-P and a recurrent novel fusion gene, *KIR2DS4/3DL1*, in HG02630-M and NA19240-P. Moreover, we identified 26 KIR alleles with variants in the CDS and 428 KIR alleles with non-CDS variants. Of the 26 alleles with CDS variants, seven were detected as synonymous with existing IPD-KIR alleles, 17 were detected as nonsynonymous, and three were detected as fusion genes composed of segments of known KIR alleles (Fig. 2e and Supplemental Table 3). Five haplotypes had ten novel alleles, 75 harbored at least five novel alleles, and all 94 haplotypes contained at least two novel alleles (Fig. 2f and Supplemental Table 2). Importantly, by annotating IPD-KIR alleles to 94 HPRC haplotypes, 115 CDS-only alleles were complemented with the full genomic sequences derived from HPRC assemblies.

### Duplications and deletions of *KIR3DP1* and *KIR2DL4* framework genes

We identified duplications of *KIR3DP1* and *KIR2DL4* in six (6.38%) haplotypes: HG002-M, HG00733-P, HG01123-P, HG02622-P, HG02630-P, and HG02717-M. These haplotypes contained two sequential copies of the *KIR3DP1-KIR2DL4-KIR3DL1/S1* triad with an intergenic length of nearly 14 Kb between *KIR3DP1* and *KIR2DL4* (Figs. 2 and 3a-b). Two consecutive triads, previously reported as tA01-ins3/4 (Pyo et al. 2013), were formed by special recombination events, and typically included a specific downstream KIR3DP1*004 allele (Pyo et al. 2013), carried by HG002-M, HG00733-P, and HG01123-P. In an earlier study, the KIR3DP1*004 allele was confirmed to be transcribed in peripheral blood mononuclear cells because of the nature of the recombinant *KIR3DP1* and *KIR2L5A* (Gómez-Lozano et al. 2005).

In contrast, deletions in *KIR3DP1* and *KIR2DL4* framework genes were observed in six (6.38%) haplotypes: HG00741-P, HG01243-P, HG01243-M, HG01358-M, HG03098-P, and NA19240-M. HG00741-P resulted from a deletion event involving *KIR2DL1-KIR3DP1-KIR2DL4-KIR3DS1-KIR2DL5A-2DS3S5* in the cA01-tB01 haplotype, which generated the cA01-tB01-del7 haplotype (Pyo et al. 2013) (Pyo et al. 2013) (Figs. 2b and 3d). The remaining five (5.32%) haplotypes, HG01243-P, HG01243-M, HG01358-M, HG03098-P, and NA19240-M, were formed by the deletion of *KIR2DP1-KIR2DL1-KIR3DP1-KIR2DL4-KIR3DS1-KIR2DL5A-2DS3S5* in the cB01-tB01 haplotype, creating the cB01-tB01-del7 haplotype (Pyo et al. 2013) (Figs. 2d and 3c).

### Full and partial deletion of *KIR3DL2*

Previous studies have reported haplotypes lacking *KIR3DL2* or exhibiting a partial deletion of *KIR3DL2* in which sequences from intron 3 to exon 9 are missing (Jiang et al. 2012; Pyo et al. 2013). In our analysis, HG00741-P exhibited a partially deleted *KIR3DL2* with a sequence matching only exons 1–3 of KIR3DL2*00601 (equivalent to exons 1–3 of KIR3DL2*10002 in our computational assignment) (Fig. 3d). This sequence was identical to the coding regions of exons 1–3 of KIR3DL2*007, as reported previously (Norman et al. 2009; Traherne et al. 2010). We also identified HG03540-M, the daughter of a Gambian trio that lacks the *KIR3DL2* framework gene and carries KIR3DL1*05901. This allele type is frequently observed in Africa and has been reported as a fusion gene of *KIR3DL1* and *KIR3DL2* resulting from a deletion event in *KIR3DL1* exons 6–9, *KIR2DS4* and *KIR3DL2* exons 1–5 within the tA01 haplotype, forming the tA01-del9-hybd1 haplotype (Norman et al. 2009; Pyo et al. 2013) (Fig. 3e).

Real-time quantitative polymerase chain reaction (qPCR) was used to further verify CNVs in the KIR framework genes in 11 HPRC samples (HG002, HG00733, HG00741, HG01243, HG01358, HG02622, HG02630, HG02717, HG03098, HG03540, and NA19240) (Supplemental Table 4) (Jiang et al. 2016). First, we confirmed that all 11 samples had two copies of the *KIR3DL3* framework gene compared to the internal control gene *STAT6*. Five samples (HG002, HG00733, HG02622, HG02630, and HG02717) carried three copies of *KIR3DP1* and *KIR2DL4*; four samples (HG00741, HG01358, HG03098, and HG03540) had only one copy of the pair; and one sample (HG01243) lacked both genes. Regarding *KIR3DL2*, a CNV with one copy was detected in two samples (HG00741 and HG03540) of the 11 studied. Because the HG00741 paternal haplotype contained only a part of *KIR3DL2* (exons 1–3), it was not detectable by qPCR. The determination of CNV in the KIR framework genes was consistent with the results of the SKIRT pipeline.

### Identification of a novel fusion gene, *KIR2DS4/3DL1*

We identified two copies of *KIR2DS4* in HG02630-M and NA19240-P, both of which are cA01-tA01 haplotypes. One of the two *KIR2DS4* copies in each haplotype was a fusion of *KIR2DS4* and *KIR3DL1* genes, comprising exons 1–7 of KIR2DS4*00101 and exons 8–9 of KIR3DL1*03501. Both haplotypes contained complete versions of KIR2DS4*00101 and KIR3DL1*03501 (Fig. 4a). We verified the presence of the *KIR2DS4/3DL1* fusion gene using PacBio HiFi reads obtained from the National Center for Biotechnology Information (NCBI) Sequence Read Archive (SRA) for both HG02630 and NA19240. We found that multiple PacBio HiFi reads perfectly matched at least exon 7 to exon 9 sequences of the novel fusion gene in both samples (Supplemental Table 5). Because *KIR2DS4* was absent in the other haplotypes of HG02630 and NA19240 (i.e., HG02630-P and NA19240-M), these PacBio HiFi reads supported the existence of the *KIR2DS4/3DL1* fusion gene. We also explored the potential evolutionary relationship between *KIR2DS4*, *KIR3DL1,* and *KIR2DS4/3DL1* fusion genes. Several regions within intron 7 of *KIR2DS4* and *KIR3DL1* exhibited high sequence similarity, as indicated by the alignment tool and represented the crossover region of the fusion gene (Fig. 4b).

**Figure 4.**
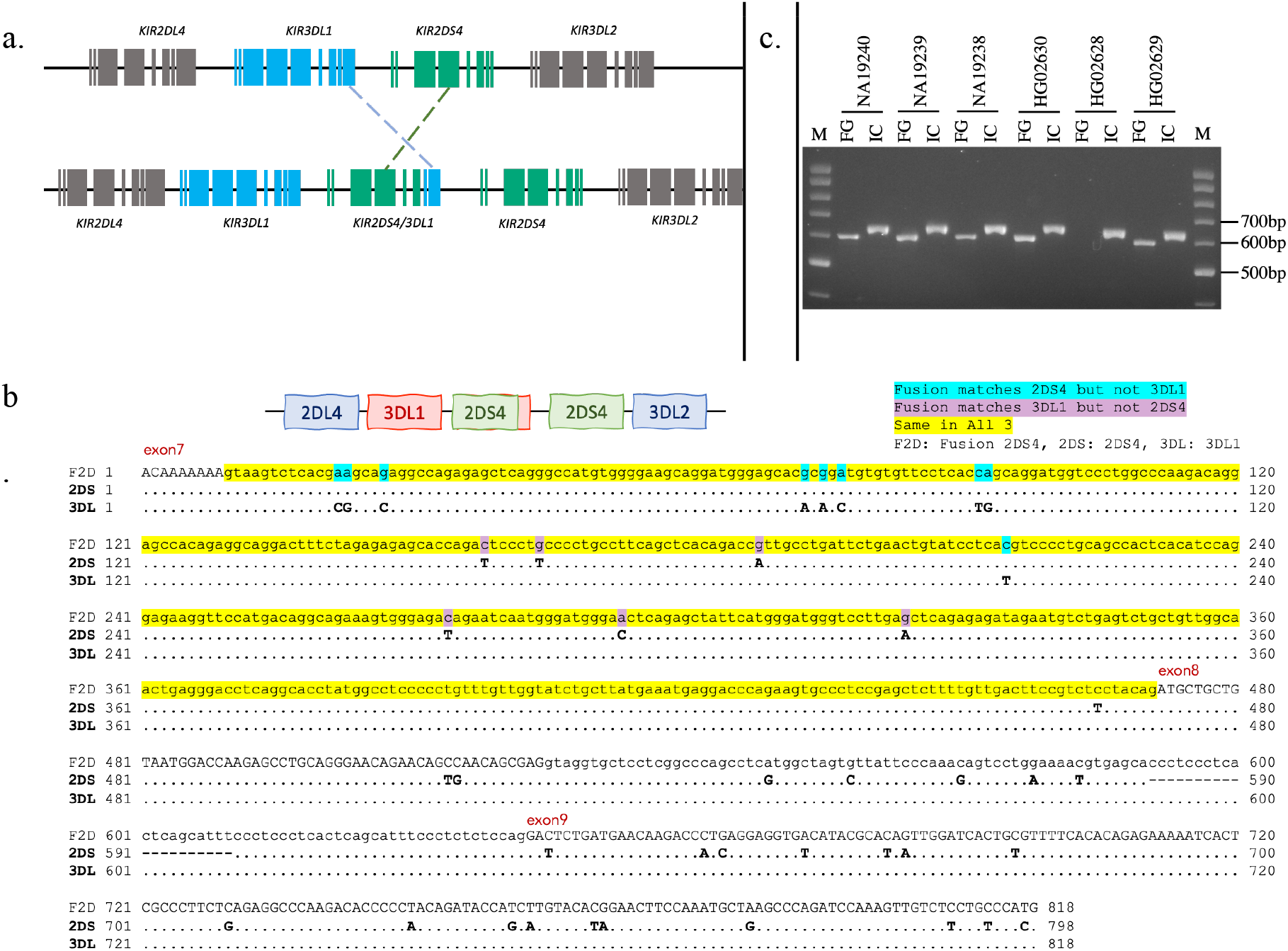
Identification of a novel fusion gene, *KIR2DS4/3DL1*. (a) A schematic diagram illustrating the formation of a novel *KIR2DS4/3DL1* fusion gene composed of exons 1–7 from *KIR2DS4* and exons 8–9 from *KIR3DL1*. (b) Sequence alignments comparing the intron 7 region of the fusion gene with corresponding regions in *KIR2DS4* and *KIR3DL1* for HG02630-M and NA19240-P. The highlighted yellow regions, signifying the sequences common to all three genes, are potential sites of the crossover event leading to the formation of the fusion gene. (c) An electrophoresis image displaying the results from the two trios of HG02630, NA19240, and their respective parents. Five out of the six samples contain the *KIR2DS4/3DL1* fusion gene (indicated by FG), whereas all six samples carry the *KIR3DL1* control gene as an internal control (indicated by IC).

We further confirmed the *KIR2DS4/3DL1* fusion gene by PCR amplification with sequence-specific primers (PCR-SSP) (Gómez-Lozano et al. 2002), followed by Sanger sequencing on the two trios of HG02630, NA19240, and their parents. Both HG02630 and her mother HG02629 carried the *KIR2DS4/3DL1* fusion gene, whereas her father HG02628 did not. Similarly, NA19240 and her father NA19239 carried the fusion gene, whereas her mother NA19238 also carried the fusion gene unexpectedly. All six samples from the two trios were positive for *KIR3DL1*, which was used as an internal control (Fig. 4c). We subsequently performed Sanger sequencing of five samples positive for the fusion gene (HG02630, HG02629, NA19240, NA19239, and NA19238) using the PCR-SSP amplicons. All fusion gene sequences in the five samples were identical to the sequences from the HPRC assemblies, confirming the presence of the *KIR2DS4/3DL1* fusion gene. Sequencing results for the *KIR3DL1* control gene showed that all five samples had identical *KIR3DL1* sequences in the amplicon region, providing evidence for the coexistence of *KIR3DL1* and *KIR2DS4/3DL1* in these samples.

### Identification of the *KIR2DL3/2DP1* fusion gene in HG02723-P

We identified the *KIR2DL3/2DP1* fusion gene in the paternal haplotype of HG02723 (HG02723-P), consistent with previous reports (Traherne et al. 2010; Hou et al. 2012; Pyo et al. 2013). In these earlier studies, the same fusion gene was detected and verified using the GenBank accession numbers CU041340 and CU633846. Our analysis of HG02723-P revealed that the fusion gene, composed of exons 1–5 of KIR2DL3*00101 and exons 6–9 of KIR2DP1*00201, matched the sequences in GenBank accessions CU041340 and CU633846.

The identification of the *KIR2DL3/2DP1* fusion gene in the KIR gene complex confirms its presence and highlights the need to investigate the potential functional implications and roles of such hybrid genes in immune responses and disease susceptibility.

### Difficulties and incompleteness in the HG002 assembly

The paternal haplotype of HG002 contained KIR3DL3*00101-KIR2DL3*00101-KIR2DP1*00201-KIR2DL1*0030218-KIR3DP1*0030202-KIR2DL4*0050101-KIR3DS1*0130101-KIR2DL5A*0010103-KIR2DS5*0020111-KIR2DS1*00201-KIR3DL2*00701 from 5-prime end to 3-prime end in one single contig. However, the HG002 maternal haplotype assembled into four different contigs containing both *KIR2DL4* and *KIR3DL1/S1*, indicating four possible copies of the KIR genes. The four contigs of the maternal haplotype were KIR3DL3*00301-KIR2DS2*005-KIR2DP1*00102-KIR2DL1*0040105-KIR3DP1*0030101-KIR2DL4*0050101-KIR3DS1*01301 in contig #1, KIR2DL4*0050101-KIR3DS1*01301 in contig #2, KIR2DL4*0050101-KIR3DS1*0130118-KIR2DL5A*00101-KIR2DS5*00201-KIR2DS1*00201-KIR3DL2*00701 in contig #3, and KIR3DS1*0130118-KIR3DP1*004-KIR2DL4*0050101 in contig #4, respectively (Fig. 5 and Supplemental Table 1). Based on the annotation, we observed that there were two complete KIR3DS1*0130118 and KIR2DL4*0050101 alleles in both contigs #3 and #4, but displayed in reverse order, whereas only partial *KIR3DL1/S1* alleles in contigs #1 and #2 were of the same order. According to the HPRC data, we could only assume that there may be more than one copy of the triad of *KIR3DP1-KIR2DL4-KIR3DS1* genes.

**Figure 5.**
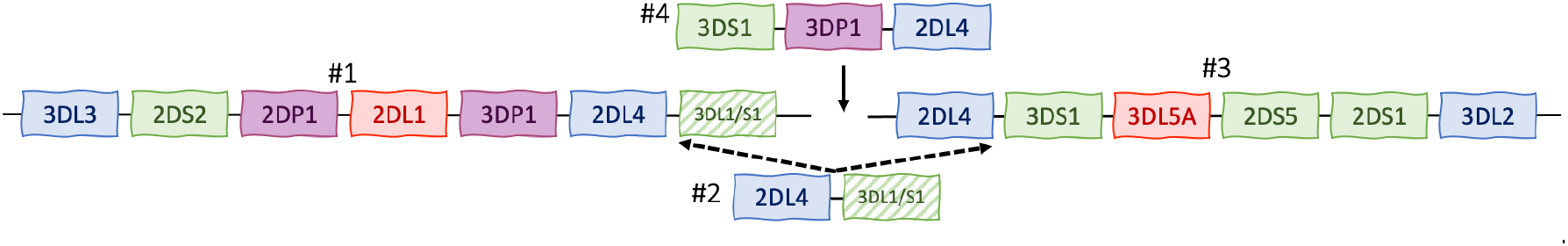
The discontinuous KIR contigs of HPRC HG002 maternal assembly. The HPRC-released HG002 maternal assembly is segmented into four contigs matching with IPDKIR alleles, representing an unusual case (1 of 94 haplotypes) in the HPRC release. Notably, no single contig encompasses the entire KIR complex. One contig (#4), *KIR3DS1-KIR3DP1-KIR2DL4*, aligns exclusively between the centromeric and telomeric motifs, suggesting potential structural variations and gene CNVs around the recombinant region.

We assessed the qualities of the KIR region in the maternal haplotype of HG002 using assemblies generated from different algorithms (Supplemental Table 6). To address the discontinuity of the HG002 maternal haplotype, we incorporated assembly data from an extended study that enhanced Hifiasm to produce more complete versions of the haplotype-resolved HG002 contig. The data, which were derived from a combination of PacBio HiFi reads and Hi-C data (Cheng et al. 2022), affirmed our primary assumption of duplication of the *KIR3DP1-KIR2DL4-KIR3DS1* triad (Figs. 2d and 3a). Specifically, the datasets named hifiasm.hic.0.16.1.rep4.hap1, hifiasm.0.16.1.bp.hap2 and hifiasm.0.16.1.primary accurately displayed two copies of the triad in a single contig, whereas HiCanu.alter displayed two copies across three distinct contigs. However, the hifiasm.trio.0.16.1.hap2 displayed three copies in one contig, whereas FALCON-PHASE.hic.hap2 had only one copy in one contig. Furthermore, we compared the KIR alleles of high-quality HG002 assemblies released in a recent study that utilized a combination of Oxford Nanopore Technologies long-read sequences, Hi-C, trio data, and/or HiFi read sequences (Rautiainen et al. 2023). We observed maximum concordance among the various datasets (downsampled_deepcns_hifiasm_hic, downsampled_deepcns_hifiasm_trio, downsampled_deepcns_verkko_hifi_ont, downsampled_deepcns_verkko_hifi_ont_hic, downsampled_deepcns_verkko_hifi_ont_trio, downsampled_verkko_hifi_ont, and downsampled_verkko_hifi_ont_trio, verkko_hifi_ont_hic), with hifiasm.hic.0.16.1.rep4.hap1, as a representative example (Figs. 2d and 3a). The only discordance was the allele of *KIR2DS5* of HG002-M, which appeared in all results of samples named starting as downsampled_verkko. Interestingly, the telomeric motifs of the paternal and maternal haplotypes of HG002 were identical, particularly in the three copies of *KIR2DL4* and *KIR3DS1*. This extreme similarity makes phasing of the two haplotypes particularly challenging. Despite the difficulties and incompleteness of HG002 assembly, the SKIRT pipeline comprehensively identified all KIR alleles of HG002 using HPRC assembly, including *KIR2DP1, KIR3DP1*, and all KIR framework genes in various copy numbers (0/1/2 copies), with no mix-ups among KIR2D genes.

### Comparisons with Kass in HG002 genome

The Kass assembly and annotation workflow did not identify the duplication of the *KIR3DP1-KIR2DL4-KIR3DS1* triad as disclosed in the study. As for HPRC HG002 assembly, Kass annotation could not generate annotations for *KIR2DP1* and *KIR3DP1* even if it perfectly matches some alleles, whereas SKIRT identified a perfectly matched allele (i.e., paternal *KIR2DP1*) and other partially matched ones. Furthermore, Kass annotation seemed to operate under the presumption of a single copy for all KIR framework genes on one haplotype and exhibited confusion regarding some KIR2D genes (Supplemental Table 1). However, SKIRT identified all the KIR alleles and CNVs of the KIR framework genes in the HPRC HG002 assembly.

### Comparisons with T1K in HPRC genomes

T1K (Song et al. 2023) offered an innovative methodology for KIR genotyping using Illumina short-read WGS, termed hereafter as T1K-genotype. The study also annotated KIR alleles in 26 of the 47 HPRC phased assemblies to validate their genotyping results, termed hereafter as T1K-annotate.

The T1K-annotate results for 26 HPRC genomes were assumed to be the benchmark for validating T1K-genotyping results, implying that the genotyping results should have significantly benefited from the annotations. However, when we used the copy number detection results of HPRC via SKIRT as the ground truth, T1K-genotype showed a recall rate of less than 90% for 13 out of 17 genes and an overall recall rate of 85% (426/499), while T1K-annotate showed a 99.6% (497/499) overall recall rate (Fig. 6a). Furthermore, the precision and recall rates for copy numbers of individual genes were largely inconsistent between T1K-genotype and T1K-annotate. T1K-annotate failed to identify the *KIR2DS/3DL1* fusion gene in the HG2630-P and the truncated *KIR3DL2* in HG00741-P, the latter of which led to incorrect identification of copy number variation of *KIR3DP1* in HG00741-P.

**Figure 6.**
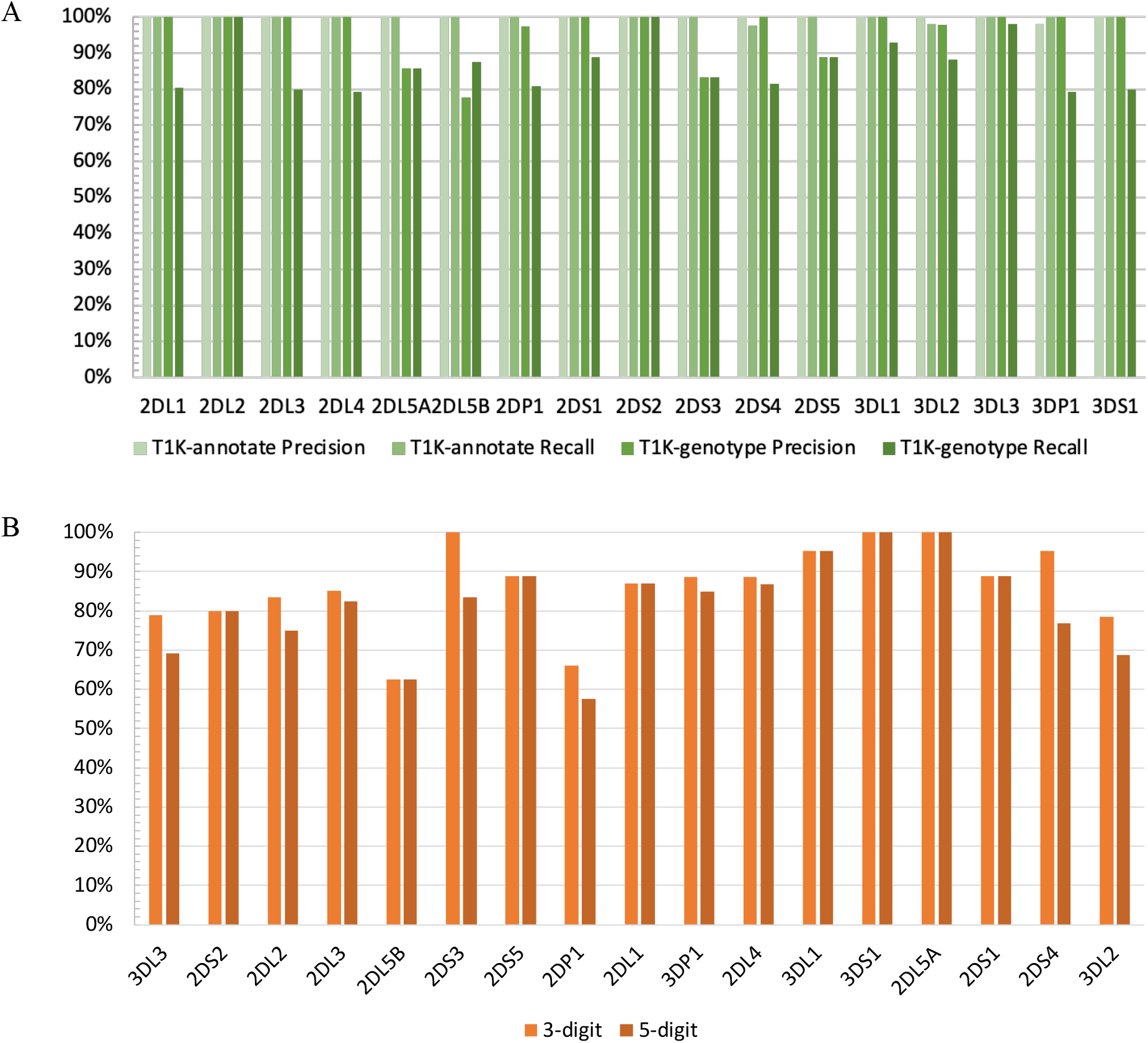
The comparisons of T1K and SKIRT. (a) The precision and recall of copy numbers determined by T1K-annotate and T1K-genotype as compared to SKIRT in 26 HPRC genomes. The gene copy numbers detected by T1K-annotate and T1K-genotype are compared separately with the results of SKIRT as the ground truth. T1K-annotate exhibits perfect precision and recall of 100% in almost all KIR genes, except for a 98% recall rate for both *KIR2DS4* and *KIR3DL2* and a 98% precision rate for *KIR3DP1*. Nevertheless, T1K-genotype shows a recall rate of less than 90% for 13 out of 17 genes. (b) The agreement between T1K-annotate and SKIRT for the allele annotation of 26 HPRC genomes. Each allele annotation of T1K is compared with the results of SKIRT at two resolution levels, 3-digit and 5-digit. If a candidate allele annotated by T1K matches that of SKIRT in the same haplotype, it is marked as one in agreement. If T1K gives one or multiple candidate alleles with differences only in the sixth and seventh digits compared to the one assigned by SKIRT, the agreement was assigned on 5-digit level. If the first three digits were the same but the fourth and fifth digits were different, the agreement was assigned on 3-digit level.

While T1K-annotate demonstrated proficiency in identifying the copy numbers of KIR genes, its performance in allele identification showed an overall agreement of 85% and 79% with the 3-digit and 5-digit resolution results from SKIRT, respectively (Fig. 6b, Supplemental Table 7). We further validated the identified alleles from both T1K and SKIRT by visually inspecting the alignments with IPD-KIR allele coding sequences, which confirmed the accuracy of the annotations made by SKIRT (Supplemental Figs. 1a-b). Given that T1K-annotate selects alleles based solely on the alignment of exonic sequences to HPRC assemblies using BWA-MEM (Song et al. 2023), we confined our comparison of allele resolutions to the coding sequence, representing the first five digits. In contrast, SKIRT provides a more comprehensive genomic resolution, extending to all seven digits.

### KIR alleles of Human Reference Genomes

We compared the SKIRT KIR gene annotations for GRCh37 (hg19) (Church et al. 2011), GRCh38 (hg38) (Schneider et al. 2017), and T2T CHM13v2.0 (hs1) (Nurk et al. 2022) with the latest GENCODE annotations available on the UCSC Genome Browser (Kent et al. 2002; Raney et al. 2014; Nassar et al. 2023) for the three reference genomes (Fiddes et al. 2018; Frankish et al. 2021; Shumate et al. 2021) (Supplemental Figs. 2a-c and Supplemental Tables 8-10). SKIRT annotations were added as custom tracks on the UCSC Genome Browser for comparison with the GENCODE annotations in the KIR region. Notably, SKIRT provided IPD-KIR allele annotations for the genomes and only displayed coding regions, excluding untranslated regions (UTRs), as no UTR information was available for the IPD-KIR allele sequences. The annotations for *KIR2DP1* and *KIR3DP1* pseudogenes were more comprehensive: exon 3 of *KIR2DP1*, identified as a pseudo-exon, was not displayed in our annotations but was shown in the GENCODE annotations, whereas exon 3 of *KIR3DP1*, which is not a pseudo-exon, was displayed in our annotations but missed in the GENCODE annotations. GENCODE sometimes provides multiple alternative splicing annotations for a single KIR gene; however, our annotations identified the exact correct coding regions for each KIR gene.

### Incomplete alleles in the IPD-KIR database and our genomic identification

In our study, we uncovered that several alleles listed as ‘Partial’ in the IPD-KIR database, regardless of their categorization as CDS-only or genomic, had either portions (KIR2DS4*00104 and KIR2DS5*009) or entirety (KIR3DL2*024, KIR3DL2*01101, and KIR3DL3*030) of exons 1 and 2 missing from their IPD-KIR listings. Our comprehensive investigation led to complete genomic identification of these alleles. Furthermore, we identified the exon 2 presence of two additional allele sequences, KIR3DP1*00601 and KIR3DP1*01001, which were confirmed as “full genomic” but without exon 2 and its flanking region in the IPD-KIR database. The KIR3DP1*00601 allele from HG01123-M diverged from the KIR3DP1*0060101 allele in the IPD-KIR database, primarily by the inclusion of an additional exon-2 segment and its flanking region, with the others of the CDS remaining identical. Similarly, the KIR3DP1*01001 allele from NA21309-P mainly differed from the KIR3DP1*0100102 allele in the IPD-KIR database in the presence of an additional exon-2 segment and its flanking region, whereas the others of the CDS were completely identical. Previous studies have indicated that certain *KIR3DP1* alleles show variations in the presence or absence of exon 2 (Gómez-Lozano et al. 2005; Bono et al. 2018).

## Discussion

This study presented a comprehensive investigation of the KIR complex in HPRC Year-1 assemblies, efficiently describing the high-resolution allele-level haplotypic structure of each chromosome pair with all existing KIR gene duplications, deletions, and CNVs. We also reported the discovery of a novel fusion gene, *KIR2DS4/3DL1*, in different populations. The SKIRT pipeline is a practical tool for analyzing KIR genes in human genome assembly data, enabling accurate identification and annotation of KIR alleles and their structural variations.

The findings of this study demonstrate that the copy numbers of the three KIR framework genes, except for *KIR3DL3*, presumably one for each haplotype in some previous studies using NGS, were mostly one but sporadically two or none for a single haplotype of the HPRC samples studied. The identification of the *KIR2DS4/3DL1* fusion gene in HG02630 and NA19240 individuals and subsequent PCR verification highlighted the importance of using a robust analytical pipeline to uncover novel KIR gene structural variations. This is particularly relevant considering the potential presence of undiscovered KIR receptors resulting from continuously occurring unequal crossover and rearrangement of gene segments in the KIR complex. The duplication and deletion of KIR genes may alter their expression and thus modulate the reciprocal immune response, such as *KIR3DL1*/*S1* influences some infectious diseases (Boudreau et al. 2016; Boelen et al. 2018; Li et al. 2022).

Utilizing the highest-resolution and accurate annotation of KIR alleles, our study effectively identified all known KIR alleles listed in the IPD-KIR database. Moreover, we provided full genomic sequences for known CDS-only alleles, as well as novel synonymous and nonsynonymous alleles. In total, we generated 570 full-genomic sequence alleles, of which 26 alleles exhibited CDS variants, 428 alleles displayed non-CDS variants, and an additional 116 alleles showed no variants but complemented their corresponding CDS-only alleles in the IPD-KIR database. This achievement paves the way for the submission of these sequences to the database, thereby catalyzing further KIR studies. Correct submission to the IPD-KIR database is critical for the advancement of KIR research. For instance, our study first identified KIR2DL5*0020102 allele for HG01952-P, KIR2DL5*0020106 for HG02145-M, and KIR2DL5*0020104 for HG02257-P. Currently, these three allele sequences are available only for CDS, instead of full genomic sequences, in the IPD-KIR database. This discovery underscores the importance of diligent and meticulous analyses in KIR research.

The HPRC Year-1 release provides data from multiple sequencing technologies, allowing researchers to access genome assemblies constructed using PacBio HiFi long-read sequences, and phased using parental Illumina short-read sequences. The raw data for these long and short sequences, obtained from publicly available cell line samples from the National Human Genome Research Institute (NHGRI) Sample Repository for Human Genetics Research at the Coriell Institute, were selected from 1KGP, which offers diverse human genetic variations (The 1000 Genomes Project Consortium 2015). These cell line samples containing DNA serve as invaluable resources for further experiments and assays, and the variety of sequencing data facilitates the development of novel analysis methods. Although there have been many studies focusing on 1KGP, T1K and SKIRT were the only three generating KIR alleles using 1KGP genomes and SKIRT outperformed the other two (Roe et al. 2020; Song et al. 2023). The high-resolution and accurate KIR alleles derived from the HPRC cohort in our study offer invaluable resources, empowering researchers to develop more comprehensive assays and advanced computational methods tailored for sequencing KIR genes. Anticipating the near future, we envision the generation of an ever-expanding repertoire of high-quality personal assemblies through the efforts of HPRC and other genomic projects. As this landscape evolves, SKIRT will continue to serve as an indispensable asset, benefiting the global KIR community.

Our results revealed several specific patterns of KIR gene duplication within the haplotypes. The most noticeable was duplication of the *KIR3DP1-KIR2DL4-KIR3DS1* triad. Through deep and high-resolution analysis of 47 high-quality genome assemblies from the HPRC, we deciphered more structural variations in the KIR complex, especially in the region close to the recombinant hotspot between the two framework genes, *KIR3DP1* and *KIR2DL4*. Long-range deletion and duplication of sequence segments in the KIR region creates a great variety of copy numbers and polymorphic alleles, constituting the deletion and duplication of KIR genes, such as *KIR3DP1*, *KIR2DL4*, *KIR3DL1/S1*, and *KIR3DL2*, and occasionally the fusion of novel genes like *KIR2DS4/3DL1* or *KIR2DL3/2DP1*.

These recombination events may be the main origins of these structural variations. The sequence similarities were so high that mapping between allele sequences and assemblies was sometimes very challenging for assigning an accurate allele. The well-organized KIR nomenclature synergizes with the IPD-KIR database (Marsh et al. 2003), curating hundreds of nucleotide and peptide sequences and facilitating the digital identification of KIR allele sequences, ideally to benefit our study. Investigating the impact of KIR CNVs and structural variations on the repertoire and function of NK cells could provide insights into the mechanisms underlying KIR gene-associated disease susceptibility and resistance. Expanding the analysis of KIR gene diversity to include a broader range of populations could help elucidate population-specific patterns of KIR gene variations and their potential implications for disease risk and transplantation outcomes.

Based on the discovery of accurate and high-resolution alleles of all existing KIR genes in these high-quality assemblies of HPRC samples, our study could promote future exploration of more effective and efficient tools and functional assays targeting the identified novel alleles and genes. In contrast, the identification of KIR alleles may be highly affected by the quality of the contigs around the KIR region. Therefore, the SKIRT pipeline could be an effective tool for evaluating the quality of assembled contigs.

## Methods

### HPRC assembly data collection and IPD-KIR database

The SKIRT pipeline annotates publicly available HRPC genome assemblies with all official KIR alleles assigned by the KIR Nomenclature Committee. We used version v2.12.0 of the IPD-KIR hosted by the European Molecular Biology Laboratory’s European Bioinformatics Institute (EMBL-EBI) (https://www.ebi.ac.uk/ipd/kir/). We obtained 47 HPRC genome assemblies from Year-1 Sequencing data. The 47 high-quality haplotype-resolved genome assemblies were produced as diploid contigs using parental Illumina short-read sequence data to sort filial haplotypes of PacBio HiFi long-read sequence data and de novo assembly using the graph trio binning algorithm of Hifiasm (Cheng et al. 2021), providing the highest resolution for haplotype-resolved contigs. With a long sequencing length (10–20 Kb) and highly accurate consensus quality (Q30) of HiFi-reads, the reads could cover each KIR gene (4–16 Kb) or intergenic region (2.4–14 Kb). Consequently, phased diploid genome assemblies and algorithms offer better ways to identify each parental KIR haplotype.

### SKIRT - Algorithms of the Structural KIR annoTator Pipeline

The pipeline was designed to identify new variants, novel alleles (both synonymous and nonsynonymous), and structural variations, including gene duplications and deletions. (Fig. 1a) To discern multiple polymorphic alleles at each potential locus, we employed minimap2 (Li 2018), a versatile pairwise long-read sequence aligner, to map all KIR alleles (including CDS-only and genomic) as query sequences against human genome assemblies as target sequences. A separate mapping process was conducted to detect the short (8 bp) exon 9 of *KIR3DS1* in the CDS-only queries, employing the ‘--end-seed-pen 5’ setting to overcome the minimap2 limitation for “tiny exons.” To detect long intragenic insertions, we activated the splice mode and limited the insertion size with the ‘-G16k’ setting. This approach facilitated the recognition of splicing sites and intragenic distances of CDS-only alleles and prevented the potential overextension issue of a single CDS-only allele mapping across multiple KIR loci due to high polymorphism among certain alleles of different genes.

Candidate aligned alleles were assessed based on several criteria for each KIR gene, including a reasonable flanking region with an intergenic length of 2.4 Kb among most KIR genes and 14 Kb between *KIR3DP1* and *KIR2DL4*, full genomic length of each gene ranging from 4 to 16 Kb, starting and ending coordinates, and numbers of mismatches and gaps. We compared all mapped candidates to select the best matching KIR alleles, achieving up to seven digits of KIR allele nomenclature indicative of genomic resolution (Fig. 1b). The mappings of genomic alleles were examined only when the same CDS-only allele perfectly matched the locus. Allele resolution was presented as seven-digit when the genomic allele matched perfectly. In the case of any variance in the mapping of the query CDS-only allele, we identified the CDS of the KIR locus in the target assemblies using the cs tag of each alignment generated by minimap2. The target CDS was mapped against the IPD-KIR protein sequences using tblastn (Camacho et al. 2009) and Biopython (Cock et al. 2009) to confirm whether the genetic variances were synonymous with an existing allele (Fig. 2e and Supplemental Table 3).

The SKIRT generates a <SAMPLE name>.hap.csv file for an overview of the entire haplotype, displaying all high-resolution KIR alleles in the order of the chromosome. However, it generates VCF and BED files, capturing allele annotations and their variants compared to the most similar alleles identified during the process. The assembly FASTA, VCF, and BED files were loaded into the Integrative Genomics Viewer (IGV) (Robinson et al. 2011) for a graphical view of the KIR allele annotations. This visualization facilitated the understanding of relative gene locations, coordinates, and structural variations, such as CNVs, deletions, gene fusions, and variants specified in the BED files (Supplemental Fig. 3).

### Identification of the fusion genes, *KIR2DS4*/*3DL1* and *KIR2DL3*/*2DP1*

Initially, we only identified a part of KIR2DS4*00101 as the *KIR2DS4*/*3DL1* fusion gene in both the maternal haplotype of HG2630 and the paternal haplotype of NA19240 because the allele mapped 96.1% of the gene segment. The predominant part, encompassing only exons 1 to 7 of KIR2DS4*00101, exhibited the absence of the subsequent exons 8 and 9. This is primarily because of the mapping preference of minimap2, which tends to identify the more extensive mapping when one locus maps to two allele sequences. In such cases, minimap2 does not present the mapping of the minor part, that is, *KIR3DL1*, in the two haplotypes, showing only those mapped to the majority of the loci. Further investigation of the downstream flanking sequences of this partial *KIR2DS4* revealed the presence of two other exon sequences completely identical to exons 8 and 9 of KIR3DL1*03501 in the HG2630-M and NA19240-P contigs. Without long and continuous genomic sequences exceeding 16 Kb in such regions, it would not have been possible to identify the fusion genes and determine the accurate copy numbers of the related genes in the haplotype, that is, *KIR2DS4* and *KIR3DL1*. We verified the existence of the fusion gene sequences using both Illumina WGS short reads and PacBio HiFi long reads, confirming our results.

We identified the *KIR2DL3/2DP1* fusion gene in the paternal haplotype of HG02723 (HG02723-P), consistent with previous reports (Traherne et al. 2010; Hou et al. 2012; Pyo et al. 2013). In these earlier studies, the same fusion gene was detected, verified, and submitted to GenBank (accession numbers: CU041340 and CU633846). Our analysis of HG02723-P revealed that the fusion gene, composed of exons 1–5 of KIR2DL3*00101 and exons 6–9 of KIR2DP1*00201, matched the coding sequences of GenBank accessions CU041340 and CU633846. This fusion gene was not a novel finding. However, the allele sequence was not submitted to IPD-KIR. We used Illumina WGS short reads and PacBio HiFi long reads to certify the sequence of this fusion gene of HG02723-P and confirmed the existence of the fusion gene at the locus. The *KIR2DL3/2DP1* fusion gene was first discovered in a sample of European ancestry (Traherne et al. 2010) and was reported again in a sample of Asian ancestry (Hou et al. 2012), both as centromeric A and telomeric B haplotypes. However, HG02723-P is a Gambian strain with the centromeric A and telomeric A haplotypes.

### TaqMan multiplex real-time PCR for CNV verification

For KIR copy number determination, we conducted TaqMan multiplex real-time PCR on an Applied Biosystems (ABI) QuantStudio™ 5 Real Time Detection System (ThermoFisher Scientific, USA) using the qKAT protocol (Jiang et al. 2016). For the KIR framework genes *KIR3DL3, KIR3DP1, KIR2DL4, KIR3DL2,* and the internal control gene *STAT6*, a reaction volume of 20 μL contained a final concentration of 1X qPCRBIO Probe Mix Lo-ROX (PCR BIOSYSTEMS), optimized concentrations of primers and probes (Supplemental Tables 11 and 12), and eight ng of genomic DNA. To achieve optimal amplification performance, *KIR3DL3* and *KIR3DP1* were detected in standard real-time PCR mode (95 °C for 3 min, 35 cycles of 95 °C for 15 s and 64 °C for 45 s), and *KIR3DL3*, *KIR2DL4*, and *KIR3DL2* were in fast mode (95 °C for 3 min, 35 cycles of 95 °C for 5 s, 65 °C for 20 s). All assays were multiplexed with the internal control gene STAT6 (Supplemental Table 13). The comparative Ct method (delta-delta Ct method, ΔΔCt) was used to determine the copy numbers of *KIR3DP1*, *KIR2DL4*, and *KIR3DL2*. The ΔCt was calculated by the Ct value of the KIR gene minus that of STAT6, then the copy number of two for the *KIR3DL3* gene was confirmed. The ΔΔCt value was the ΔCt of a particular KIR gene (*KIR3DP1*, *KIR2DL4*, or *KIR3DL2*) minus the ΔCt of *KIR3DL3* to determine its copy number.

### PCR-SSP primer design and confirmation of *KIR2DS4/3DL1* fusion gene

To amplify the correct product of the *KIR2DS4/3DL1* fusion gene, primers were designed for PCR-SSP (Gómez-Lozano et al. 2002), specifically for HG02630 and NA19240. Our design ensured that the primer sequences perfectly matched the specific target alleles, whereas others had certain nucleotide mismatches, particularly at the 3’ end of the primers. We examined the exon7 sequence of *KIR2DS4* to determine the forward primer, KIR2DS4_fusion, and the exon 8 sequence of *KIR3DL1* to obtain the reverse primer, KIR3DL1_reverse. These primers were designed to produce PCR amplicons 596 bp in length. We also designed another forward primer targeting on the exon7 of *KIR3DL1*, KIR3DL1_control, to ensure the effectiveness of the primer sets. Together with the KIR3DL1_reverse primer, this primer set produced 634-bp amplicons when *KIR3DL1* gene was present in the samples (Table 1).

**Table 1.**
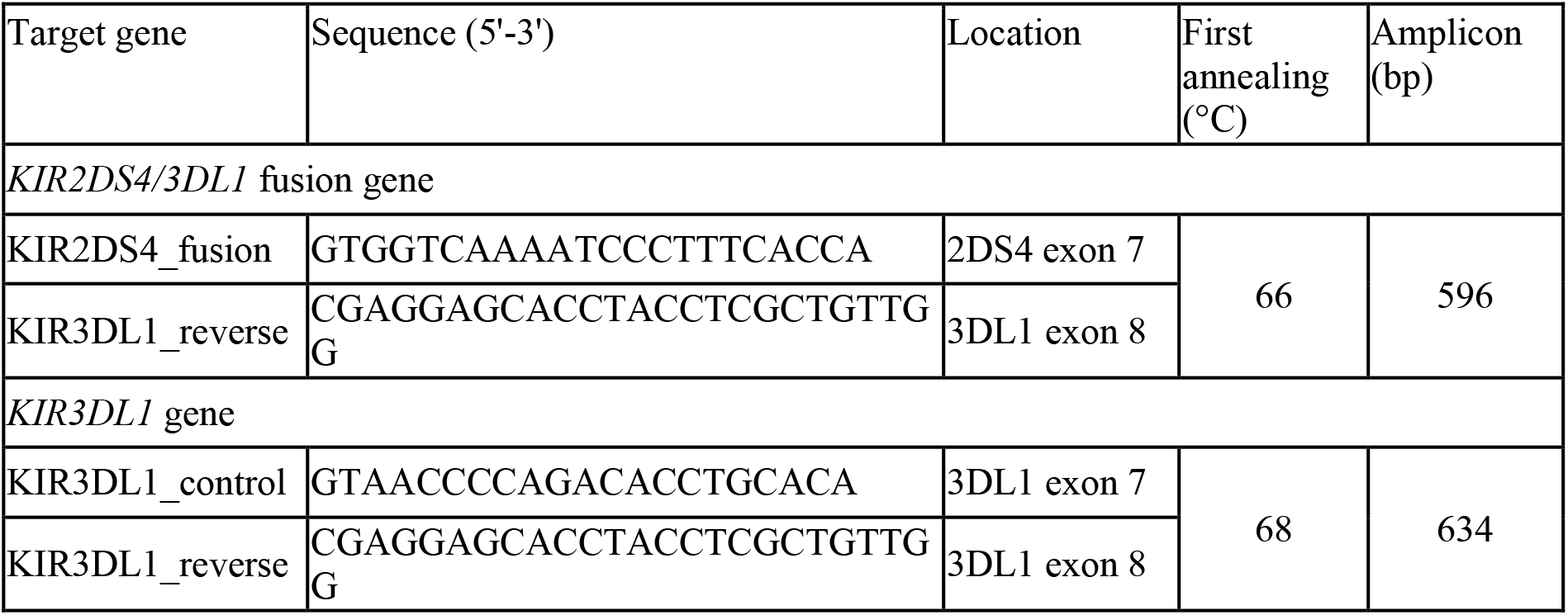
Primer design for the amplification of *KIR2DS4/3DL1* fusion gene.

Two observed samples and their parents (<u>HG02630</u>, HG02629, HG02628, <u>NA19240</u>, NA19239, and NA19238) were processed using two-step PCR assays on the Thermocycler TProfessional Standard 96 Gradient (Biometra®) system. The reaction volume of 50 μL contained a final concentration of 1X GoTaq® Green Master Mix (Promega), 400 nM of each forward and reverse primers and 150 ng genomic DNA. The PCR process was hot-start at 96°C for 5 min and then the first amplification of 15 cycles at 96 °C for 30 s, an annealing temperature of 66 or 68 °C (fusion gene and the control gene, respectively) for 30 s, and 72 °C for 1 min, followed by 25 cycles at 96 °C for 30 s, 64 °C for 30 s, 72 °C for 1 min, and final extension at 72 °C for 10 min. After amplification, the PCR products were analyzed using 4% agarose gel electrophoresis.

Sanger sequencing of the PCR products was conducted to confirm the presence of the *KIR2DS4/3DL1* fusion gene and *KIR3DL1* control gene in five samples (HG02629, HG02630, NA19238, NA19239, and NA19240) that showed positive results in both PCR assays.

## Data accesss

All HPRC cell lines were obtained from Coriell, with the exception of HG01123, HG02486, and HG02559. The PacBio HiFi data used in this study were obtained from NCBI Sequence Read Archive (SRA) under the BioProject accession number PRJNA701308 for HG00411, HG00423, HG00438, HG00480, HG00621, HG00673, HG00730, HG00735, HG00741, HG01071, HG01106, HG01123, HG01138, HG01175, HG01258, HG01358, HG01361, HG01888, HG01891, HG01928, HG01952, HG01978, HG01998, HG02027, HG02083, HG02148, HG02257, HG02486, HG02523, HG02559, HG02572, HG02622, HG02630, HG02717, HG02886, HG03453, HG03471, HG03516, HG03540, HG03579, and PRJNA731524 for HG002, HG005, HG00733, HG01109, HG01243, HG01442, HG02055, HG02080, HG02109, HG02145, HG02723, HG02818, HG02970, HG03098, HG03486, HG03492, NA18906, NA19030, NA19240, NA20129, NA20300, and NA21309. The NCBI SRA accession PRJNA730823 is an umbrella project for all HPRC-related projects and PRJNA730822 is an umbrella project for all 47 HPRC assembly projects. The other two assembly datasets for HG002 are available publicly at Zenodo (https://doi.org/10.5281/zenodo.5948487; https://doi.org/10.5281/zenodo.7400747) (Cheng et al. 2022; Rautiainen et al. 2022). All data generated or analyzed during this study are included in this published article, its Supplemental files and Zenodo (https://doi.org/10.5281/zenodo.8094803). SKIRT source code is publicly available on GitHub (https://github.com/calvinckhung/skirt). All genomic sequences of novel KIR alleles except KIR2DP1 and KIR3DP1 were submitted to the NCBI GenBank with submission identifiers. Accession numbers of NCBI for KIR2DL1 alleles were obtained (BK064711-BK064753), while other KIR genes are awaited and will be obtained (Supplemental Table 14).

## Competing interest statement

The authors declare that they have no competing interests.

## Acknowledgments

This study was funded by the Ministry of Science and Technology, Taiwan (MOST-109-2622-B-002-004-CC2). We thank the National Center for High-performance Computing (NCHC) of National Applied Research Laboratories (NARLabs) in Taiwan for providing computational and storage resources. The English editing of this article was sponsored by National Taiwan University with the support of Higher Education Sprout Project from the Ministry of Education, Taiwan.

## Author contributions

P-L.C., J.S.H., Y-C.Y. and Y-C.L proposed the initial idea of annotating KIR alleles with high-quality assemblies, initiated the project, and assisted with discussions.

T-K.H. conceived the study, designed the analysis pipeline, acquired the HPRC data, performed the analysis, interpreted the analysis results, and produced the related figures and tables with P-L.C.’s, Y-C.Y.’s, J.S.H.’s, and C-L.H.’s guidance.

W-C.L. and Y-C.Y. designed all laboratory experiments, and W-C.L. performed the experiments and produced related figures and tables under the guidance of Y-C.Y..

T-K.H. and W-C.L. managed the writing and organization of the manuscript with suggestions and proofreading by Y-C.Y and J.S.H.

H-W.C. and H-Y.L. assisted with the WGS data access and analysis under the guidance of J.S.H. and C-Y.C.

P-L.C. managed the project.

All authors read and approved the final manuscript.

## Notes

### Competing Interest Statement

The authors have declared no competing interest.

